# Lipid remodeling by hypoxia aggravates migratory potential in pancreatic cancer while maintaining membrane homeostasis

**DOI:** 10.1101/2022.12.08.519694

**Authors:** Prema Kumari Agarwala, Shuai Nie, Gavin E. Reid, Shobhna Kapoor

## Abstract

Membranes are crucial cell components underlying optimal cellular functioning under diverse conditions including cancer. The membrane physiology requires acute maintenance of biophysical properties and a regulation of cellular lipidome. Homeostatic adaptation of membranes to temperature, pressure and anti-cancer drugs is a well-recognized. However, how the same is regulated under the influence of oxygen deprivation in pancreatic cancers-highly hypoxic cancer- is not known. Here, we report robust lipidomic remodelling in response to HIF-1α induction in pancreatic cancer cells and significant accumulation of lipid droplets. The lipidome rewiring span changes across various lipid classes, levels of unsaturation and acyl chain lengths. Interestingly, despite extensive lipidome alteration, cellular membrane homeostatic response ensures no major modulation of membrane biophysical properties underlying enhanced migratory potential. The correlation of lipidome changes, with pathway analysis and proteomics provide the basis for mutually exclusive regulation of lipidome and membrane properties. These findings help to understand the hypoxic regulation of pancreatic membrane homeostasis.

## Introduction

Pancreatic cancer (PaC) is a highly aggressive malignancy, with a less than 9% five-year survival rate and poor prognosis [1]. The dismal prognosis of PaC is due to its rapid disease progression and highly metastatic potential [2]. At a genomic level, a mutation in KRAS is the key step underlying tumour initiation, followed by mutations of TP53, and p53 driving malignancy towards the metastatic tumour [3]. In addition to genetic and epigenetic alterations, pancreatic cancer also involves significant alterations of cellular metabolism including that of glucose, amino acids, fatty acids (FA), and lipids, indispensable for their growth and survival [4, 5]. Further, multi-omic profiling of pancreatic cancer stem cells shows extensive reliance on the fatty acid pathway for survival implying exquisite remodeling of pancreatic cancer lipidome[6]. Another major hallmark of PC is the presence of oxygen deprivation, i.e., hypoxia, which is correlated strongly to cancer aggressiveness [7-9]. Hypoxia induces the stabilization of hypoxia-inducible factor 1α (HIF-1α), which dimerizes with HIF-1β, transfers into the nucleus, and binds with hypoxia-responsive elements present in DNA [10]. Hypoxic gene expression downstream of HIF-1α is thought to be a major driving force in many cancer progressions coupled with robust rewiring of many cellular metabolic pathways. Specifically, hypoxic rewiring of lipid metabolism has revealed a tight regulation of lipid synthesis, lipolysis, and lipid droplets [10]. Extracellular FA flux, TG synthesis, and synthesis of derivative phospholipids increase under hypoxia, while lipid catabolism is suppressed. Notably, the accumulation of cholesterol has also been reported [11]. To obviate lipotoxicity due to an imbalance between synthesis and catabolism, lipid storage in lipid droplets is aptly enhanced [12]. The latter serves as the major source for the production of essential sterol esters and importantly phospholipids for the biogenesis of new membranes.

Modulation of lipidome foremost affects membranes which are essential for the physiological functioning of mammalian cells, by maintaining the cellular structural integrity. Furthermore, non-structural roles of lipids e.g. signaling, protein function, membrane fusion, and trafficking have also emerged [13, 14]. Membranes orchestrate various membrane-associated processes by leveraging their biophysical properties such as fluidity, packing, stiffness, curvature, and lateral organization [15]. Changes in these properties alter lipid/protein diffusion, localization, lipid-protein interactions, and finally their activity [16, 17]. These impose an exquisite control on regulating cellular processes and hence are critically regulated in cancer for cell survival and proliferation [18]. For instance, fluidity in cancer is altered both ways, while in highly metastatic cancer, the fluidity is increased by reduction of cholesterol and saturated lipids for facile movement (deformable membranes), in drug-resistant cancer, the fluidity is decreased to restrict permeability to drugs [18]. Likewise, cell cancer membrane mechanics and lateral organization are linked with their metastatic dissemination. Collectively, because of a central role in cellular functioning, effective maintenance of membrane properties is essential and, hence the homeostatic response of mammalian membranes is crucial in both normal and disease states.

The correlation between the inherently high hypoxic stress in pancreatic cancer, cellular lipidome remodeling and subsequently membrane properties and metastatic potential is not known. Here we directly evaluate the pancreatic cell membrane response to hypoxia by characterizing the lipidomic, biophysical, and cellular alterations. Using mass spectrometry we show distinctive lipidome changes in cancer cells versus normal cells under the influence of HIF-1α. Levels of glycerophospholipids and sphingolipids were significantly downregulated accompanied by a disproportioned increase in the levels of cholesterol esters and triacylglycerides contributing to increased lipid storage. Distinct changes in the levels of saturation and lipid acyl chain length variations in a cell-line dependent fashion was furnished. Employing an arcade of membrane methods, we reveal decoupling of cell membrane biophysical properties from the lipidome alterations, indicative of a stable homeostatic membrane response under hypoxia. Finally, we show that this homeostatic membrane response is essential for cellular fitness under hypoxia to maintain high metastatic potential. These findings will help gain a deepened understanding of the underlying mechanisms as well as molecular pathways involved in the hypoxic regulation of pancreatic membrane homeostasis. This is a fundamental step towards the development of reliable lipid and membrane biophysical biomarkers as well as membrane-centric therapeutic interventions.

## Materials and Methods

### Materials

Foetal bovine serum (FBS, RM1112), Dulbecco’s Modified Eagle Medium (DMEM-AT007), 1X Dulbecco’s phosphate buffer saline (D-PBS), 20X Antibiotic-antimycotic solution, 0.25% trypsin-EDTA, MTT reagent and RIPA buffer were procured from HiMedia, India. Laurdan, D-5030 media, puromycin, hEGF, Cobalt (II) chloride hexahydrate (C8661) and Bradford reagent were purchased from Sigma Aldrich, St. Louis, Missouri, USA. The probes DPH (diphenylhexatriene) and TMA-DPH (trimethylamino-diphenylhexatriene) were purchased from Cayman Chemicals, USA. Di-4-ANEPPDHQ RDHPE, TRITC-Phalloidin and Bodipy dyes were purchased from Thermo Fischer Scientific. Western blotting substrate and protease inhibitor tablets were also from Thermo Fischer Scientific, USA. PhosphoSTOP tablets were purchased from Roche, Switzerland. Mouse monoclonal anti-HIF1α antibody was purchased from BD Biosciences, San Jose, CA, Mouse monoclonal anti-β-actin antibody from LI-COR Biosciences and HRP-conjugated goat monoclonal anti-mouse secondary antibody was from Thermo Fischer Scientific. M3 base medium without antibiotic was purchased from INCELL Corporation, LLC.

### Cell culture

PANC-1 and CAPAN-2 pancreatic ductal adenocarcinoma (PDAC) cells were provided as a generous gift by Prof. Ritu Aneja from the Georgia State University, Atlanta. The normal HPNE cell line was purchased from ATCC. PANC-1 & CAPAN-2 cell lines were cultured in DMEM medium and HPNE cells were maintained in D5030 medium supplemented with M3 base medium.

### Cobalt Chloride (Cocl_2_) treatment to mimic hypoxia

Cocl_2_ was dissolved in autoclaved MilliQ water and the stock was diluted in serum free medium (0.4%FBS) to achieve the required concentration.

### Cell viability assay

Cytotoxic effect of Cocl_2_ was evaluated using MTT assay. Briefly, all the pancreatic cells were seeded in a flat bottomed 96-well polystyrene coated plate at a seeding density of 1*10^4^ cells in duplicates. After 36 h cells were treated with different concentrations of Cocl_2_ from 800 µM to 50 µM (2 fold dilutions) for 24 h in high and low serum media. After treatment with each diluted compound and incubation for 24 h, 100µL of MTT (0.5 mg/ ml) was added to each well and incubated for 3 h at 37°C. Formazan crystals formed after 3 h in each well were dissolved in 150 μL of DMSO and the plates were read immediately in a microplate reader (BIO-RAD microplate reader-550) at 570 nm. Wells without cells were used as blanks. This method was used to determine the cell viability.

### Western blot analysis

To check the HIF1-α protein expression, cells were seeded on 60-mm plates and incubated for 36 h. After the treatment with 200 µM Cocl_2_ in low and high serum medium for 24 h, cells were scraped off, washed once with ice-cold phosphate-buffered saline (PBS) and then lysed using RIPA buffer supplemented with 0.1% beta mercaptoethanol, protease and phosphatase inhibitor cocktails. Samples were incubated on ice for 1 h with in between vortexing for 30 sec after each 5-10 min and centrifuged at 14000 RPM for 30 min. The total protein concentration was measured using Bradford method. Samples were heated at 95 °C for 10 min with 1X SDS loading buffer and briefly cooled at room temperature and then 50-60 μg of the total proteins from each sample were resolved to 10% SDS-polyacrylamide gel electrophoresis (PAGE) to detect HIF-1α (120 kDa) and β-actin (42 kDa). Resolved proteins were then electrophoretically transferred onto polyvinylidene difluoride (PVDF) membranes (Merck Millipore, USA) at 90 V for 2.5 h, and then blotted membranes were blocked with 5% skim milk in 1X TBST for 1.5 h at room temperature. Blotted proteins were probed with the corresponding primary and secondary (HRP conjugated) antibodies. Pierce™ ECL western blotting substrate was used to detect the bands on a LI-COR gel imaging system. With β-actin as a control, the relative expression of each protein was calculated for each experimental group.

### Lipid extraction and liquid chromatography - mass spectrometry (LC-MS) analysis

After aspiration of the media, cultured cells were washed with PBS and quenched with 0.75 mL of ice cold methanol/PBS (1:1, v/v). The cells were then scraped into tubes, sonicated, followed by addition of SPLASH lipidomix internal standard mix (330707, Avanti Polar Lipids) and deuterated ceramide lipidomix internal standard (330713X, Avanti Polar Lipids) at 10 μL per 1*10^6^ cells. This was vortexed for 1 min, followed by addition of 0.5 mL chloroform, vortexed for 2 min and centrifuged at 500 g for 10 min. Finally, the chloroform layer was transferred to a new glass vial, dried under nitrogen gas and stored at -20 °C. Immediately prior to LC-MS analysis, samples were reconstituted in 50 µL of chloroform/methanol (1:1 v/v).

Lipid analysis was performed using a Q Exactive Orbitrap^TM^ mass spectrometer (Thermo Scientific). The LC parameters were as follows: 3 µL of sample was injected onto a 1.7 µm particle 100 *2.1mm ID Waters Acquity CSH C18 column (186005297, Waters) which was kept at 50 °C. A gradient of (A) water/acetonitrile (40:60, v/v) with 10 mM ammonium acetate and (B) acetonitrile/2-propanol (10:90, v/v) with 10 mM ammonium acetate at a flow rate of 0.3 mL/min was used. The gradient ran from 0% to 40% B over 6 min, then from 40% to 100% B in the next 29 min, followed by 100% B for 4 min, and then returned to 0% B in 2 min, where it was kept for 4 min (45 min total). Lipids were identified in both positive and negative modes. The electrospray and mass spec settings were as follows: spray voltage 3.2 kV (positive mode) and 3.6 kV (negative mode), capillary temperature 320^0^C, sheath gas flow 50 (arbitrary units) for negative mode and 45 (arbitrary units) for positive mode and auxiliary gas flow 8 (arbitrary units) for negative mode and 12 (arbitrary units) for positive mode. The mass spec analysis was performed in a full MS and data dependent MS2 (Top 5) mode, with a full scan range of 300-1000 m/z, resolution 70,000, automatic gain control at 1x10^6^ and a maximum injection time of 250 ms. MS2 parameters were: m/z scan range of 200-2000, resolution 17, 500, automatic gain control was set at 1x10^5^ with a maximum injection time of 200 ms, dynamic exclusion 6S and the normalized collision energy (NCE) was as follows 20,35,50 [19].

LC-MS/MS data for lipidomic study was searched through MS Dial 4.90. The mass accuracy settings were 0.005 Da and 0.025 Da for MS1 and MS2. The minimum peak height was 5000 and mass slice width was 0.05 Da. The identification score cut off was 80%. Post identification was done with a text file containing name and m/z of each standard (SPLASH and Deuterated Ceramide lipidomix Mass Spec Standard). In positive mode, [M+H]+, [M+NH4]+ and [M+H-H2O]+ were selected as ion forms. In negative mode, [M-H]- and [M+CH3COO]-were selected as ion forms. All lipid classes available were selected for the search. The retention time tolerance for alignment was 0.1 min. Lipids with maximum intensity less than 5-fold of average intensity in blank was removed. All other settings were default. All lipid LC-MS features were manually inspected and re-integrated when needed. These four types of lipids, 1) lipids with only sum composition except SM, 2) lipid identification due to peak tailing, 3) retention time outliner within each lipid class, 4) LPA and PA artifacts generated by in-source fragmentation of LPS and PS were also removed. The shorthand notation used for lipid classification and structural representation follows the nomenclature proposed previously [20].

Relative quantification of lipid species was achieved by comparison of the LC peak areas of identified lipids against those of the corresponding internal lipid standards in the same lipid class and their concentrations at nmol/L reported by the supplier. Finally, the lipid species at the class, subclass or molecular species levels were normalized to either the total lipid concentration (i.e., mol% total lipid), or total lipid-class concentration (i.e., mol% total lipid class). For the lipid classes without correspondent stable isotope-labelled lipid standards, the LC peak areas of individual molecular species within these classes were normalised as follows: the MG species against the DG (18:1D7_15:0); the LPG against the PG(18:1D7_15:0), the LPA against the PA(18:1D7_15:0) and the LPS against the PS (18:1D7_15:0).

Given that the commercially available stable isotope-labelled lipid standards are limited, some of the identified lipids were normalised against a standard from a different class or sub-class, and no attempts were made to quantitatively correct for different ESI responses of individual lipids due to concentration, acyl chain length, degree of unsaturation, or matrix effects caused by differences in chromatographic retention times compared with the relevant standards. The results reported here are for relative quantification and should not be considered to reflect the absolute concentrations of each lipid or lipid sub-class.The normalized data were then used for statistical analysis. Significantly dysregulated lipids annotated using volcano plots were determined using Metaboanalyst 5.0.

### Pathway enrichment analysis using Lipidsig

Lipid related gene enrichment analysis was performed using web based lipid sig tool. The database associated with this tool is KEGG and the method used was Fisher’s exact test. The top N significant terms and gene similarity were kept at 30 and 0.5 respectively. The *P* value cut off was kept at 0.05 [21].

### Sample preparation for proteomic analysis

Peptide preparation from protein cell lysate:

Pancreatic cells were seeded and after the treatment were subjected to lysis and protein extraction using a urea lysis buffer containing 8 M urea, 50 mM Tris pH 8.0, 75 mM NaCl, and 1 mM MgCl_2_ with phosphostop and protease inhibitor cocktail. After sonication, cell lysates were then centrifuged at 20,000 *g* for 20 min, followed by quantification of protein concentration of the clear lysates using Bradford assay. 50µg of protein lysates were subsequently reduced with tris (2-carboxyethyl) phosphine (TCEP) at 37°C for an hour, and alkylated using iodoacetamide for 10 min in the dark. Samples were diluted 5-fold with 25 mM Tris pH 8.0 and 1 mM CaCl_2_ prior to digesting them with trypsin (Promega, V511X) at a 1:30 enzyme-to-protein ratio at 37 °C in a dry bath for 16 h.

Digested peptides were acidified with formic acid (FA; Fluka, 56302) to a final volumetric concentration of 1% or final pH of ∼3–5, and centrifuged at 2000 *g* for 5 min to clear precipitated urea from peptide lysates. Samples were desalted on (C18-sep packs column, Merck Milipore), vacuum-dried and dissolved in 0.1% FA.

Liquid Chromatography-Tandem Mass Spectrometry (LC-MS/MS):

The MS system was equipped with an automated Easy-nLC 1200 system coupled to LC-MS/MS (Orbitrap fusion) and a linear gradient of solvent A (0.1% FA in water) to solvent B (0.1% FA, 80% ACN) was run for 120 min in positive mode. The analytical C18 column (ES903A, 50 CM) used was 50 cm in length, 75 µm in diameter and particle size 2 µm. The flow rate was kept at 0.3 µl/min. Ion source type kept was NSI and ion transfer tube temperature was at 320°C. The spray voltage and gas mode was in static mode. The voltage for positive and negative ion was 2100 and 600 respectively. MS OT mode was set with a resolution of 60000, scan range of 375-1800 m/z, maximum injection time of 50 ms, mass tolerance of 10 ppm with a dynamic exclusion duration of 60 sec and ddMS² IT HCD having a resolution of 15000, isolation window 1.6, with a maximum injection time of 30 ms [22].

Label free proteomics data analysis by proteome discoverer 2.4:

The raw data sets were processed using proteome discoverer (PD; version 2.4. Using the parameters discussed in a previous study [23]. In brief, the variable modification of carbamidomethyl and oxidation was set as static modification. The search engines used were MASCOT and SEQUEST with the Homo sapiens database. False-Discovery Rates (FDRs) for peptide and protein identifications were set to 1%. For data normalization the ‘Total peptide intensity’ parameter was used.

### Steady-State Fluorescence Anisotropy

For the study of membrane fluidity by steady state fluorescence anisotropy, 3*10^5^ of each cells were seeded per well in 6 well plate and allowed to attach for 36 h. Cells were treated for 24 h with 200 µM Cocl_2_ in low serum condition. Post treatment, cells were washed with 1X PBS, trypsinized, centrifuged for 5 min at 2500 rpm, and suspended in 1 mL of PBS. Laurdan, TMADPH and DPH probes were dissolved in DMSO and added to achieve a final probe concentration of 5 μM and 4 μM, respectively. Cells were incubated at room temperature for 20 min, centrifuged at 2500 rpm for 10 min and re-suspended in PBS. Fluorescence anisotropy was measured using a Varian Cary fluorescence spectrophotometer with a temperature controller equipped with Varian manual polarizer accessory (Australia). The cell suspension after transferring to a cuvette was kept sometime to attain temperature of 37°C using a circulating water bath. The samples were excited with vertically and horizontally polarized lights and respective polarized emission intensities were recorded. The degree of fluorescence steady-state anisotropy (*r*) was calculated from the following equation:

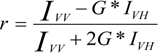

where *I_VV_* and *I_VH_* are the vertically polarized emission intensities measured in directions parallel (both at 0°, *I*_0_) and perpendicular (excitation: 0°, emission: 90°, I_90_) to the excitation beam. The correction factor (G= *I_HV_/I_HH_*) is the ratio of the detection system sensitivity for the polarized light, where *I_HV_* and *I_HH_* corresponds to the emission intensities perpendicular and parallel to the horizontally excitation light. The correction due to cell scattering was applied for obtaining the final anisotropy values [24].

### Laurdan GP Spectroscopy

Laurdan, a fluorescence solvatochromic probe, is sensitive to the solvent polarity was employed to determine the membrane hydration and order [25, 26]. The emission spectra of Laurdan exhibit spectral shifts due to dielectric relaxation which can be analyzed by calculating generalized polarization (*GP*). All the steps for sample preparations were same as the sample preparation of steady state fluorescence anisotropy experiments. Laurdan was excited at 350 nm and the emission was measured from 380 nm to 550 nm. The emission spectra of Laurdan were recorded in pancreatic cancer cells in presence and absence of Cocl_2,_ using the Varian Cary Eclipse spectrofluorometer and the shift in the spectrum of Laurdan were quantified using the Generalized Polarization function which can be defined as the following:

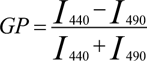

*I_440_* and *I_490_* refer to the intensities at 440 nm and 490 nm, which are characteristic for an ordered (gel) lipid phase and fluid liquid-crystalline phase, respectively. The correction due to cell scattering was also applied to obtain the final *GP* values coming from the probe. The *GP* values ranges from +1 to –1.

### Membrane labelling with Di-4ANEPPDHQ and Confocal Spectral Imaging

Pancreatic cells were seeded on glass bottom dishes and after the treatment washed with 1XPBS, followed by addition of 5 μM of Di-4ANEPPDHQ dye for 30 min. Prior to imaging, cells were again washed, re-suspended with fresh media and imaged on a laser scanning confocal microscope (Carl Zeiss, Germany) with excitation at 488 nm with Argon ion lasers as an excitation source. The spectral images were recorded from 490 nm to 695 nm with 10 nm intervals using 63 x oil immersion objective lens (1.4 NA). The resulting stacks of spectral images were processed and analysed using a MATLAB Spectral Imaging Toolbox [27]. Images were thresholded using intensity threshold present in the software and objects of interest (i.e., plasma membrane) were segmented using a watershed-based plugin approach. Generalized polarization (GP) was then calculated at each pixel using the intensities from the images collected at two emission maxima 584 nm (λ_Lo_) and 610 nm ((λ_Ld_) of the probe, using the following equation:

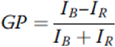

Where, *I _B_*= 584 nm and *I _R_*= 610 nm

The calculated GP values are then visualized using a pseudo-colored GP map with a look-up table scaled from -1(high fluidity) to +1.0 (lowest fluidity, ordered phase) Finally, the distribution of *GP* values is fitted to a one peak Gaussian distribution and the resultant *GP* histogram can then be used to calculate changes in lipid order or disorder.

### Atomic Force Microscopy

Force spectroscopy was done in contact mode with MFP-3D atomic force microscope (Asylum Research, Santa Barbara, CA, USA.). Silicon nitride cantilevers with spring constant 37.55 pN/nm was used and 60nN force was applied; velocity of cantilever was kept constant at 5 µm/s. Force curves were recorded only at the periphery of the cells. On each cell, at least 2 force curves were recorded. Force curve was recorded on at least 70-80 cells. For determination of elastic modulus by fitting the force curve using Hertz model, Igor software (Asylum research) was used after providing necessary inputs for tip geometry and Poisson’s ratio of the sample. For analysis of membrane tethers (tether force, tether number and lengths), a Matlab programme (designed by Peter Nagy) and freely available online was used [28].

For force spectroscopy experiments, pancreatic cells were seeded at a seeding density of 3*10^5^ at 37°C, after the treatment for 24 h cells were washed with 1X PBS, re-suspended in 3 ml of fresh serum free media and force spectroscopy of control and treated cells were recorded.

### FliptR FLIM imaging of pancreatic cancer cells

The FliptR probe was used to measure membrane tension in live cells [29]. FLIM imaging was performed using a Nikon Eclipse Ti A1R microscope equipped with a time-correlated single-photon counting module from PicoQuant58. Excitation was performed using a pulsed 485 nm laser operating at 20 MHz, and the emission signal was collected through a 600/50 nm bandpass filter using a gated PMA hybrid 40 detector and a TimeHarp 260 PICO board (PicoQuant). SymPhoTime 64 software (PicoQuant) was then used to fit fluorescence decay data (from full images or regions of interest) to a dual exponential model.

### Actin staining for Iso-surface Imaging

After the treatment for 24 h cells, pancreatic cells were fixed with 4% paraformaldehyde for 20 min at room temperature and rinsed with 1X PBS for three times. Then, TRITC-Phalloidin (1:500) was used to stain F-actin for 1 h in the dark at room temperature, rinsed thoroughly three times with 1X PBS, and slides were mounted with mounting media. Images were captured on a Laser scanning confocal microscope (Carl Zeiss 780) using a 63X/1.4NA objective. Imaris (Bitplane, Zurich, Switzerland) software was used to generate the 3D reconstruction (iso-surface images) of the filaments from the stacks of confocal images. In a total, approximately 60 to 80 cells from randomly selected ROI were analysed per condition per experiment. The volume enclosed by the iso-surfaces and the area enclosed by maximum intensity projections from confocal z-sections of cells, were then exported in excel, normalized and plotted in Graph Pad Prism 5.0, which provides a reliable quantitative estimation of the cellular F-actin content [30].

### Fluorescence Recovery after Photobleaching (FRAP)

Pancreatic cells were seeded on a glass bottom dishes and after the treatment, they were washed with 1XPBS, followed by addition of 2 μg/mL of N-Rh-DHPE dye for 3 min at room temperature, and imaging was performed at 37 °C in a humidified 5% CO_2_ atmosphere, using confocal microscopy with fluorescence recovery after photobleaching (FRAP) module. Prior to imaging the regions for bleaching, reference and background was selected. A bleaching area of 12 x 12 (circular region) pixels, which is equal to approximately (2 μm)^2^ was used, and was bleached within the acquisition interval of 2 s, considering the pixel dwell time of 1.58 μs. For continuous confocal scanning, 63x/1.4 NA oil objective was used with the confocal pinhole set at 1–2 Airy units. Bidirectional scanning was done with a frame mode of 512 x 512 pixels at 2x optical zoom. The entire experiment consisted of 500 cycles, and each cycle was run with a scan speed of 242.04 msec. Photobleaching of N-RH-DHPE was performed with a 561 nm laser and the iterations was kept at 30 with 100% laser power transmission. Pre- and post-bleach images were monitored at very low illumination laser power (2 mW).

For analysis, bleached and reference intensities were subtracted with the background intensity. To measure the recovery kinetics of fluorescent light intensity, post-bleached image series were normalized (double and full-scale normalization) and single exponential association algorithm was then used to fit the curve in Graph Pad Prism 5.0. The half-lives, diffusion coefficient (*D)* and % of mobile fraction were then plotted using the extracted values from the fitting.

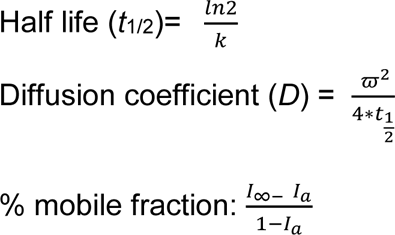

Where, I_∞_ is the full-scale normalized fluorescence intensity after full recovery and *I*_α_ the normalized intensity of the first post-bleach.

### Lipid Droplet staining with Bodipy

After the hypoxia induction for 24 h, pancreatic cells were fixed with 4% paraformaldehyde for 20 min at room temperature and rinsed with 1X PBS for three times. Cells were then incubated with 1 mM of BODIPY for 15 min in the dark, washed 2x with 1 PBS, incubated with 1 mg/mL DAPI for 15 min in the dark and washed 2x again with PBS. Coverslips were then mounted on glass slides and Z stack images were captured using confocal laser scanning microscope and processed with ImageJ software.

### Transwell migration and invasion assays

Transwell migration and invasion assays were performed to test migratory and invasive potential of pancreatic cancer cells upon hypoxia induction. Briefly, 4*10^4^ cells were seeded in Boyden chambers (PET membrane, 8 μM pore size) with or without matrigel coating (Corning), in 0.4% serum media, in presence or absence of CoCl_2_. The Boyden chambers were then inserted into 24-well plates containing DMEM supplemented with 10% fetal bovine serum as chemoattractant. Cells that migrated across the membrane were imaged 24 h post treatment. Before imaging, chambers were washed in PBS, fixed in methanol, followed by staining with a 0.5% crystal violet solution. The membranes were then imaged at 4 randomly selected fields with an inverted light microscope and cells number was calculated per field of the image.

### Statistical analysis

The results of independent experiments are presented as mean or median values as indicated. Error bars represents standard error of the mean (SEM). Statistical significance was tested using unpaired two-tailed *t*-test or two way ANOVA followed by Bonferroni’s multiple comparison test as indicated. For AFM results non-parametric unpaired Mann-Whitney test was used. *P*-values < 0.05 were considered statistically significant.

## Results

### Global lipidomics of pancreatic cancer cells under hypoxia reveals rewiring of the lipid repertoire in terms of classes, chain lengths, and unsaturation index

To investigate hypoxia-induced alterations in the global lipidome of pancreatic cell lines, PANC-1 and CAPAN-2, an in-depth quantitative mass spectrometric analysis was performed. Hypoxia was induced using non-toxic concentrations of CoCl_2_ and confirmed via HIF-1α induction (Fig. S1A-B). HPNE, a non-tumorigenic pancreatic cell was used as a control. In a total of 304 lipids were identified and quantified in PANC-1, 348 in CAPAN-2 and 276 in HPNE cells, out of which only 170 (PANC-1), 84 (CAPAN-2) and only 20 (HPNE) lipid molecules belonging to four main lipid categories such as glycerophospholipids (GPLs), sphingolipids (SPs), glycerolipids (GLs), and sterol lipids (ST) were found to be significantly altered under hypoxia compare to normoxic conditions (Fig.1A). These lipids covered 12 lipid classes such as phosphatidylcholines (PC), lyso-phosphatidylcholines (LPC), phosphatidylinositol (PI), phosphatidylethanolamine (PE), lyso-phosphatidylethanolamine (LPE), phosphatidylserine (PS), phosphatidylglycerol (PG), sphingomyelin (SM), triacylglyceride (TG), cholesteryl esters (CE), ceramides (CER) and diacylglycerol (DG). Among these specifically, TG and CE (belonging to GLs and ST categories) were significantly upregulated in cancerous cell lines. GPLs and SPs were downregulated under hypoxia (Fig. 1B). The profile of TG species with variable acyl carbon lengths and saturation levels showed a significant increase under hypoxia. No such changes were observed in HPNE under treatment (Fig. S1D). Increased abundance of CE and TG levels implicate modulation of cellular lipid droplets (LDs) under hypoxia in pancreatic cancer cells as these lipids are major components within LD for lipid storage [31]. Indeed, we observed a significant induction of lipid droplets under HIF-1α upregulation in both cell lines (Fig. 1C, with the effect being most pronounced in PANC-1 following the higher accumulation of predominantly CE in these cells (Fig. 1B)).

**Figure 1.**
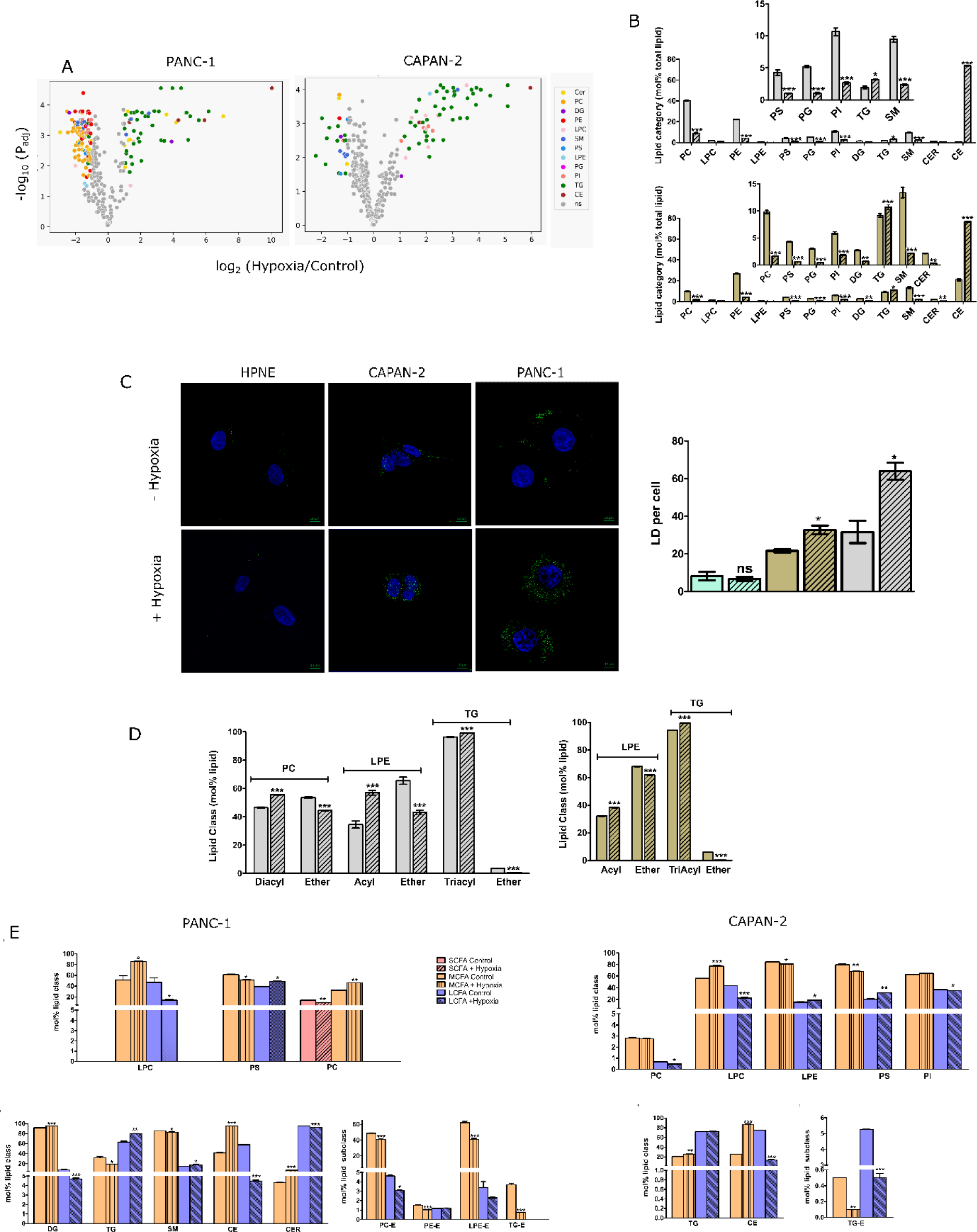
Lipidome remodeling induced by hypoxia. A. Volcano plot depicting the significant fold change of molecular lipid species at mol % total lipid in PANC-1 and CAPAN-2 cells cultured in the absence and presence of hypoxia. Lipids with a min. fold change of 2 and adjusted *p-value* < 0.05 are displayed in a color coordinated manner where blue represents lipids that are significantly downregulated and red denotes significantly upregulated lipids. B. Mol% total lipid abundance at the lipid class level shows differential distributions The inset shows the low abundance lipids for clarity. In both the cell lines, hypoxic condition down-regulates most lipid classes in glycerophospholipids (GPLs) and sphingolipids (SPs), while up-regulates most lipid classes in glycerolipids (GLs) and sterol lipids (STs). In all the cases, bar plots with patterns represents hypoxic conditions. PANC-1, CAPAN-2 and HPNE are represented by grey, brown and light green colour codes, respectively. C. Lipid droplet staining with Bodipy showed a higher number of LDs in both the cancer cells in presence of hypoxia compare to normoxia and normal cell line HPNE. Data for LDs are presented as mean ± SEM of two independent experiments and statistical analysis for LDs was determined using student’s *t*-test (\**P* < 0.01, two-tailed Student’s t-test). D. Mol% individual lipid class abundance distribution at the glycerol backbone linkage level. In both the cell lines, a significant increase in acyl-LPE and triacyl-TG was observed in hypoxic conditions, accompanied with a decrease of their ether-linkage-containing counterparts. In PANC-1 cells, diacyl PC also shows upregulation. [alkyl ether(E) = alkyl ether (O) / plasmalogen (P)]. E. Mol% total lipid class and subclass abundance to evaluate the distribution of chain lengths with individual lipid class or subclass preferred under hypoxia. A significant preference for different chain lengths is observed in a cell line-specific manner. SCFA (short carbon fatty acid), MCFA (medium carbon fatty acid), and LCFA (Long carbon fatty acid). Data represent the average mol% total lipid class abundance ± standard error of the mean of three independent experiments. Statistical significance was determined by two-way ANOVA followed by Bonferroni’s multiple comparison test. (* *P* value < 0.05, ** *P* value < 0.01, and \*\*\**P* value < 0.001).

Modulation in the mol% distribution of acyl vs. ether lipids of the most abundant lipid classes was also observed under hypoxia (Fig. 1D). For the GPLs, these included PC and LPE subclasses (i.e., diacyl, versus alkyl ether (O) and plasmalogen (P) together abbreviated as ether lipids; E (O/P)). Significant increases in mol% of diacyl-PC and acyl-LPE were observed under hypoxia in a cell-line dependent fashion. Consistent results were also reported previously in colorectal cancer [32]. Interestingly diacyl-PC only increased in PANC-1 cells. The corresponding decrease in mol% PC-E and LPE-E was observed. Again, the decrease in PC-E was limited to PANC-1 cells. A similar analysis of the GL lipid category revealed a significant increase in the mol% of triacyl-TG and a corresponding decrease in the mol% of TG-E in both cells. Collectively, these imply that in hypoxia, pancreatic cancer cells incorporate more typical acyl-containing species and downregulate the abundance of alkyl- and alkenyl-containing “ether” PC, LPE and TG lipid species, as in pancreatic normal cells a reverse trend was observed in case of LPE (Fig. S1I). This is in contrast to some reports wherein an increased abundance of ether-linked GPs is correlated to their ability to act as diagnostic markers [33]. Although, ether-containing ethanolamine phospholipids have previously been shown to undergo a significant reduction in pancreatic cancer patient serum [34]. To validate these findings in a cell-line dependent fashion, label free proteomics was performed. Reduced expression of the metabolic enzyme alkylglyceronephosphate synthase (AGPS), a critical enzyme involved in the synthesis of ether lipids including ether glycerophospholipids [35], was observed in PANC-1 proteome under hypoxia, but not in CAPAN-2 and HPNE (Fig. S2B). In addition, the abundance of AGPS in PANC-1 was higher than in CAPAN-2, accounting for a higher mol% of PC-E in normoxic PANC-1 compared to normoxic CAPAN-2 cells [36].

Next, investigating the lipid classes and subclasses based on chain length preferences revealed that hypoxia-induced an accumulation of medium chain length species within PC (C30-C39), LPC (C16-C20), DG (C30-C39), CE (C16-C20), and CER (C30-C39) in PANC-1 (Fig. 1E); LPC (C14-C20) and CE (C18) were commonly regulated in CAPAN-2 cells, and TG (C40-C50) was differentially affected. In contrast, for PS (C30-39), SM (C30-C39), and TG (C40-C50) in PANC-1, downregulation of these chain lengths was observed. Long-chained species were upregulated within PS (C40-C42), SM (C40-C52), and TG (C51-C60) lipid classes in PANC-1; only PS (C40) was similarly modulated in CAPAN-2 (Fig. 1E). In contrast, downregulation of long-chained species was observed within LPC (C22-C28), DG (C40), CE (C22), and CER (C40-C46) in PANC-1 and PC (C44-C48), LPC (C22-C28), CE (C22), and PI (C40) in CAPAN-2. Chain length variation within lipid subclasses in PANC-1 revealed that in ether-linked PC (C30-C39), PE (C33-C39), and TG (C53-C56), the medium lengths lipids were downregulated; TG-E (C46) was similarly affected in CAPAN-2. Additionally, long-chained ether-linked PC (C40-C48) was lower under hypoxia in PANC-1, while for CAPAN-2, this downregulation percolated strongly towards ether-linked TG (C52-C56) (Fig. 1D). No such variations were observed in HPNE control and treated cells except for TG medium (C36-C50) and long chains (C51-C60).

To better understand the impact of hypoxia on the saturation index of lipids, likely to impact membrane biophysical properties, we evaluated the poly-unsaturation levels in all lipid classes belonging to the four categories in all cell lines (Fig. 2A-B, and Fig. S2A). Hypoxia modulated multilevel poly-unsaturation within different lipid categories, i.e., in GPLs species with 6 double bonds; in SPs species with >6 double bonds; in GLs and STs, species with 3-4 double bonds were significantly upregulated in PANC-1 (Fig. 2A). In contrast upregulation of similar species of GLs (except for the ones with 3 double bonds) and SLs were not observed in CAPAN-2; GPLs behaved similarly and SPs did not show any significant change (Fig. 2A). Differences by lipid classes revealed that the major contributor to the observed changes in GPLs was from PS and PE in PANC-1 (Fig. 2C) and PS and LPE in CAPAN-2 (Fig. 2D). In both cell lines, increase in poly-unsaturation within PS was accompanied with an increase in corresponding chain lengths with the same species. Similarly, the major contributors for the increase of species with > 6 double bonds in SPs was mainly from SM; with concomitant increase in its chain lengths as well. For GLs and STs the changes were found to be restricted to DG, TG (along with longer chain lengths), and CE, respectively within PANC-1 (Fig. 2C). In CAPAN-2, the increase of lipid species with 3 double bonds within GLs, is contributed mainly from TGs (Fig. 2D). No similar changes were observed in HPNE (Fig. S2A).

**Figure 2.**
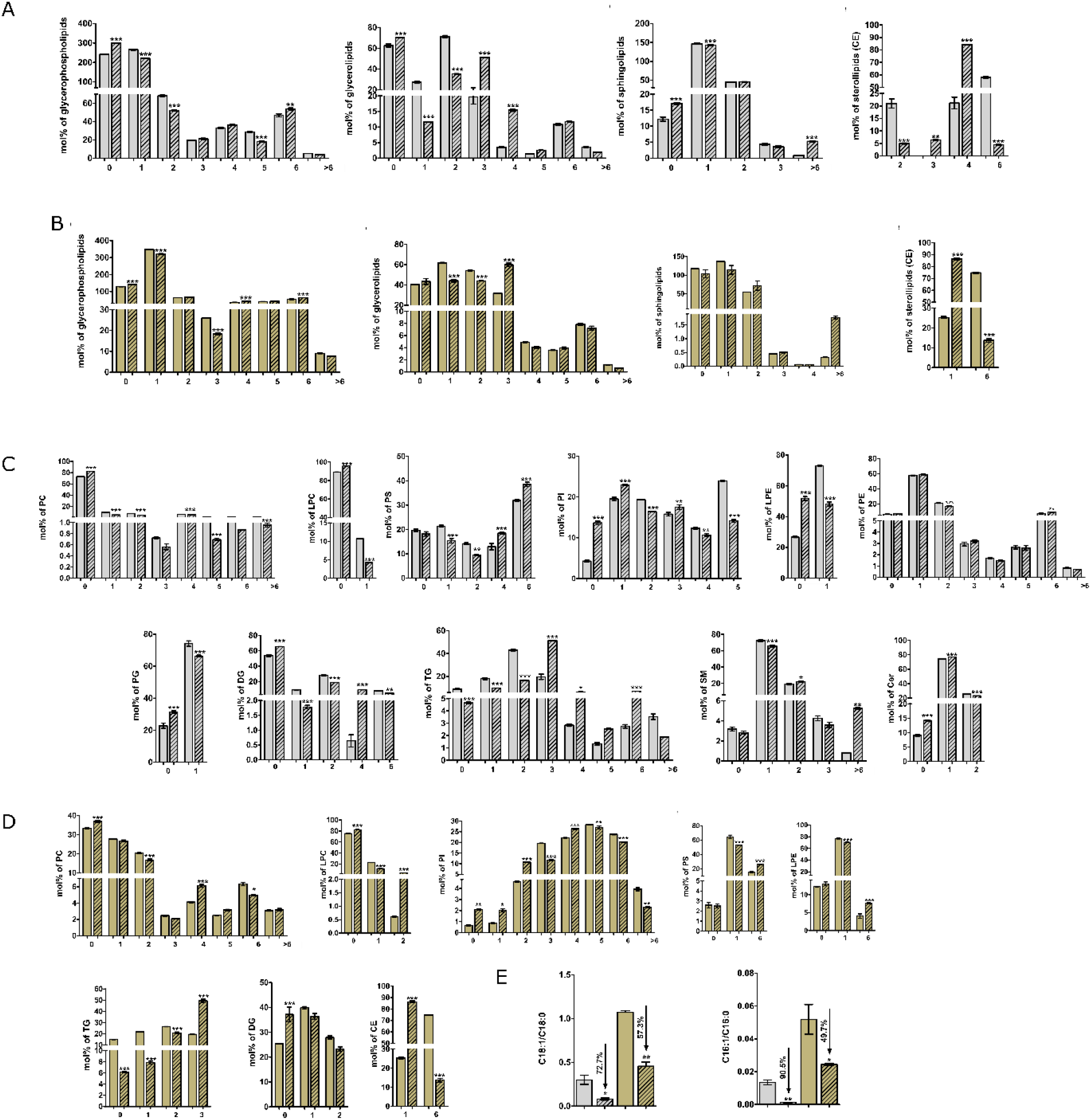
Remodeling of lipid unsaturation index at lipid category and class level.

A-D. A significantly increased unsaturation index is observed at both the lipid category (A&B) and class level (C&D) in both cell lines.

E. A lower desaturation index (C18:1/C18:0) & (C16:1/C18:0) of LPC is observed in both the cell lines.

Data represent the average mol% total lipid class abundance ± standard error of the mean of three independent experiments. Statistical significance was determined by two-way ANOVA followed by Bonferroni’s multiple comparison test. (* *P* value < 0.05, ** *P* value < 0.01, and \*\*\**P* value < 0.001).

In all the cases, bar plots with patterns represents hypoxic conditions. PANC-1, and CAPAN-2 represented by grey and brown colour codes, respectively.

Next, we evaluated the SCD1 flux or desaturation index (C18:1/C18:0) and (C16:1/C16:0). SCD1 reports on the activity of O_2_-dependent stearoyl-CoA desaturase-1 (SCD1) [37, 38]. Under hypoxia, the indices decreased in both cell lines, with the effect being most prominent for PANC-1 (Fig. 2E) supporting impaired SCD1 activity already reported under hypoxia. This also explains the increased abundances of saturated acyl chains within various lipid classes, for instance within PC, PI and LPC (Fig. 2C-D) in both the cell lines. Concomitant upregulation in levels of polyunsaturated chains in similar and different lipid classes is likely to nullify any major perturbation of the plasma membrane biophysical properties. Interestingly, the protein expression of many of the desaturases showed a complex behavior in PANC-1 cells compared to other cell lines suggesting a tight balance between saturation vs unsaturation under hypoxia. For example, the expression of FADS1 and FADS2 increased, while that of SCD5 and DEGS1 decreased (Fig. S2C). At present, it is difficult to assign which enzymes are responsible for the observed changes in the (poly) unsaturation levels of fatty acids within different lipid classes under hypoxia. In HPNE, following lipidomic profiling, no major changes were observed with these indices (Fig. S2D). Collectively, cell-line-specific changes in pancreatic cancer imply that in tumour tissues, hypoxia can induce a complicated phenotype governed by cell heterogeneity.

Our lipidomics data revealed distinct changes in the pancreatic cancer lipidome and prompted us to explore the perturbed associated lipid pathways using a web based tool, LipidSig [21]. In here, the regulated lipid classes for a particular cell type were used to get KEGG pathway enrichment analysis and all associated genes involved in that pathway. Metabolic pathways (ether lipid, inositol phosphate, glycerophospholipid, alpha-Linolenic acid, Linoleic acid and Arachidonic acid metabolism) and sphingolipid signalling pathways were ranked as the top hit in KEGG pathway enrichment analysis (Fig. 3A-B), queried with downregulated lipid classes in both PANC-1 and CAPAN-2. Endocytosis pathway was found to be the top hit for genes related to the upregulated lipid classes (Fig. 3C).

**Figure 3.**
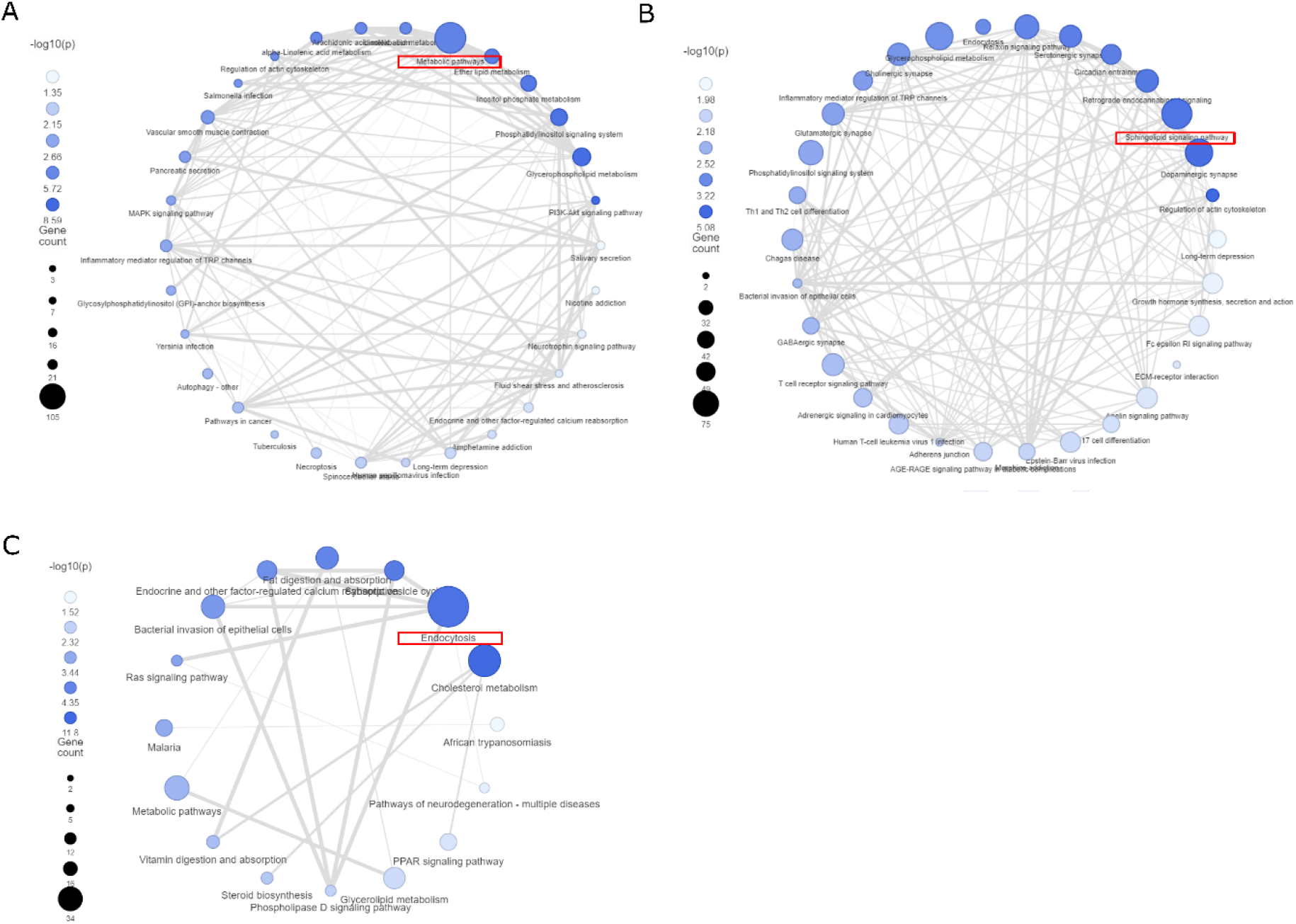
Deregulated lipid pathways in presence of hypoxia.

Enrichment network built from KEGG pathway analysis presents the significantly altered pathways (P < 0.05) associated with downregulated lipid class in A. PANC-1 and B. CAPAN-2 cells in presence of hypoxia and C. %total upregulated lipid classes in both the cell lines. Nodes are filled according to – log10 (P) and their sizes represent the lipid-related gene number involved in the pathway. Line width indicates the value of gene similarity between the pathways.

### Hypoxia-induced lipidomic perturbation preserves homeostatic maintenance of membrane biophysical properties

The lipidome remodeling induced by HIF-1α in cancer cells suggested induction of a homeostatic membrane response, wherein increases in the saturated and monounsaturated fatty acids (MUFA) chains were strongly compensated by induction of polyunsaturated fatty acids (PUFA) chains within various lipid classes in each category (Fig. 2A-D). To support this inference, we directly analyzed membrane packing in cancerous and control live cells using solvatochromic dyes (Laurdan and Di4-Aneppdhq) whose spectral characteristics are dependent on membrane properties [27, 39]. Specifically, the emission spectrum of these dyes is red-shifted in loosely packed membranes due to enhanced polarity in the vicinity of the probe within the membrane. This spectral shift can be quantified by ratio metric spectroscopy or imaging. The resulting ratio metric parameter called Generalized Polarization (GP) reports on the membrane packing and fluidity, with higher values indicative of a more tightly packed membrane. Confocal spectral imaging of Di4-aneppdhq in the live cell membranes was used to construct 2D GP maps of untreated and treated cells (Fig. 4A-B) and displayed a broad distribution of GP pixels (Fig. S3A) indicative of a heterogenous cell membranes, characteristics of mammalian cells. No effect on global membrane packing by induction of hypoxia for 24 h was seen in all cell lines (Fig. 4A-B and S3B). To gain further spatial insights, GP signals from specifically plasma membrane (PM) of live cells under normoxic and hypoxic conditions were selected and a similar behavior was observed within the PM segmented regions of the cell (Fig. 4A-B, S3B). Using the Laurdan probe and spectroscopy, similar observations confirmed the maintenance of homeostatic membrane packing and fluidity under the influence of HIF-1α (Fig. S3C). Reasons for minimal disruption to the plasma membrane could be the trade-off between saturation and poly-unsaturation levels with the plasma membrane lipids, as well as trafficking of unsaturated TG species to the plasma membrane [40].

**Figure 4.**
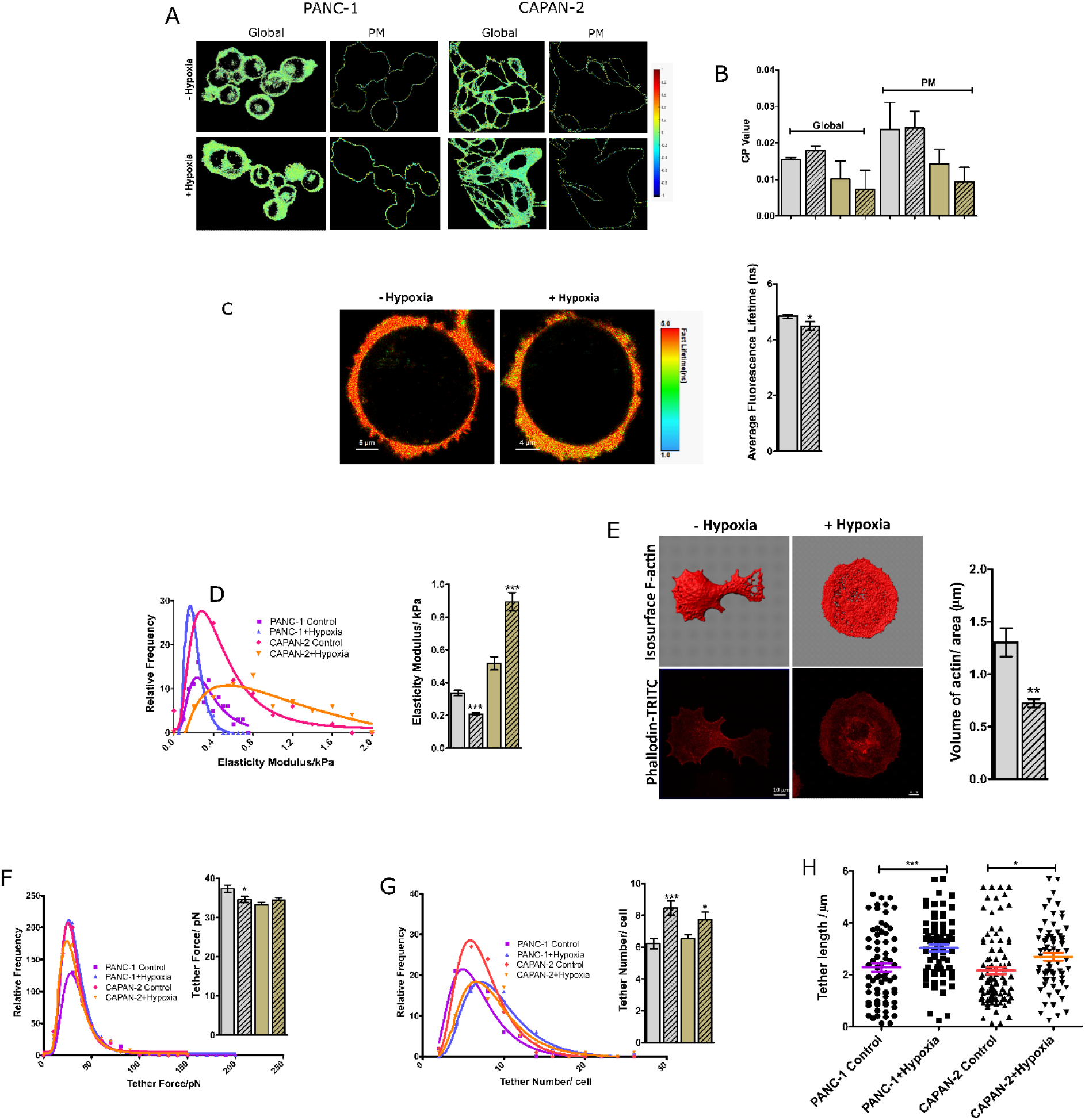
Membrane homeostasis remains intact in presence of hypoxia.

A. GP imaging of di-4-ANEPPDHQ-labeled cell membranes of both cell lines shows no change in membrane packing after 24 h of hypoxia induction.

B. GP values for both global and plasma membrane remain non-significant after the treatment in both cells.

C. CAPAN-2 cells in the presence and absence of hypoxia were used to TMADPH fluorescence anisotropy of three independent experiments. The anisotropy data showed a minor reduction in microviscosity. The statistical analysis was determined using student’s t-test (*P < 0.01, two-tailed Student’s t-test).

D. FLIM imaging of PANC-1 cells with FliptR dye showed a decrease in average fluorescence lifetime after 24 h of hypoxia induction.

E. Elastic moduli distribution of PANC-1 and CAPAN-2 cells in the presence and absence of hypoxia. A decrease in Elastic modulus values in PANC-1 and an increase is observed in CAPAN-2 cells.

F. Representative isosurface and Phalloidin-TRITC images were used to quantify actin abundance. The bar graph represents the volume of actin per unit area in control and hypoxia-induced cells. Scale bar, 10 μm; 63X oil objective.

G. Relative frequency distributions of membrane tether forces, H. tether number, and I. tether length in the presence and absence of hypoxia. The mean values are provided in the bar graph. The significance test was performed using a student’s t-test (\**P* < 0.01, two-tailed Student’s t-test).

In all the cases, bar plots with patterns represents hypoxic conditions. PANC-1, and CAPAN-2 represented by grey and brown colour codes, respectively.

Next, we evaluated lipid bilayer depth-dependent membrane fluidity/microviscosity in live cells using fluorescence anisotropy and specific lipid probes. While DPH (diphenylhexatriene) positions itself at the deep hydrophobic acyl chain regions (hydrophobic), TMA-DPH (trimethylamino-diphenylhexatriene) is located at the interfacial polar head group region of the lipid bilayers [41]. A minor decreased microviscosity of the cell plasma membrane, denoted by enhanced rotational mobility of only the TMA-DPH membrane probe was obtained in CAPAN-2 cells (Fig. 3C). This indicated a relatively fluid interfacial cell membrane region in CAPAN-2 cells compared to PANC-1 and HPNE. Cer induces membrane ordering and exhibits intermolecular hydrogen-bond capability, which increases membrane microviscosity at the interfacial bilayer regions [42]. Thus, a robust downregulation of Cer in CAPAN-2 under hypoxia is expected to alter the interfacial lipid packing as observed. Then, we analyzed membrane tension in live cell membranes under normoxic and hypoxia induction using fluorescence lifetime imaging of a mechanosensitive FliptR probe [29]. Upon hypoxia induction the average lifetime of the probe slightly decreased in PANC-1 cells while the HPNE cells depicted no change under similar conditions (Fig. 4D and Fig. S3E). Decrease of lifetime indicates the reduction in lipid phase separation, local plasma membrane composition becoming more homogenous or changes in membrane curvature [43]. A higher reduction of PC lipids; major constituents of lamellar and ‘curvature-free’ membrane in PANC-1 could account for an altered membrane tension. Collectively, no major perturbation of plasma membrane fluidity, packing and ordering under hypoxia was observed, despite extensive lipidome remodeling. Membrane dynamics were also unaffected, confirmed by measuring no substantial changes in the lateral lipid diffusion in the plasma membranes of cancer and normal live cells under hypoxic and normoxic conditions (Fig. SF-H), matching the behavior observed with control healthy HPNE cells. These suggest that HIF-1α induces lipid remodeling in hypoxic-aggressive cancers but still preserves cell membrane homeostasis for optimal membrane-associated cellular functions and survival.

Using atomic force microscopy, membrane nanomechanical properties were investigated. Hypoxia-induced cell membrane softening in PANC-1 but the reverse was observed in CAPAN-2; no change in control healthy cells (Fig. 4E and S3I). This could be explained by cell-line-dependent changes in the lipidome induced by HIF-1α. First, ether-linked lipids, which are known to modulate lipid raft domains in the plasma membrane displayed a higher downregulation in PANC-1 and hence may account for disrupted raft membrane regions and hence stiffness (Fig. 1D). Second, a higher level of poly-unsaturation with various lipid classes was observed in PANC-1, making the membrane softer (Fig. 2A and C). Third, TG species have three lipid acyl chains attached to the head group, and quantification based on the number of saturated/unsaturated chains revealed that the abundance of TGs having three unsaturated chains (i.e., 0 chains are saturated; 0 SFA) increased in PANC-1, but decreased in CAPAN-2 (Fig. S2E). Though TG is a predominant resident of lipid droplets and ER, they can also accumulate in plasma membranes (2-8 mol %) and the above findings support the nanomechanical investigation of cell membrane stiffness in a cell-line dependent fashion [40, 44]. Fourth, a softer cell membrane in PANC-1 could also stem from an increased abundance of medium chain length PC lipids, because (PC constitutes the maximum % (> 40 %) in plasma membranes (SM and PS <10-15), as the long chain lipids would enhance tighter packing of the membrane. The decreased elastic modulus in PANC-1 may also have a contribution from decreased actin filaments/volume induced by HIF-1α(Fig. 4F).

Then, we investigated the impact of hypoxia on tether dynamics in pancreatic cancer. Membrane tethers are membrane nanotubes regulating cellular communication, adhesion, and immune responses [45]. Tether force, total extension of the tethers per cell, and tether numbers indicate alterations in the membrane reservoir. Tether force is associated with membrane stiffness and bending rigidity and tether length and number indicate membrane resistance to bending. The accumulation of specific lipids can affect membrane tethers by inducing curvature and signals the plausible role of lipid sorting (induced by hypoxia during lipid remodeling) toward the formation of such curved regions at the site of insertion [28]. Hypoxia significantly reduced the median tether force in PANC-1 compared to normoxic conditions, whereas a minor increase and no change in CAPAN-2 and HPNE cells were observed, respectively (Fig. 4G and S3K). These findings support the elastic modulus’s analysis and suggest that hypoxia induction reduces the force required for the tethers to be extended, reducing surface tension and bending rigidity/elastic modulus in a cell-line-dependent fashion. (39). The tether number/cell increased in hypoxic conditions along with an increase in the total extension of the last tether (i.e. tether length) in both cancer cell lines compared to HPNE (Fig. 4H-I and S3L-M). These collectively support facile tether dynamics, which could be crucial for cancer progression under hypoxic states. Furthermore, the formation of the membrane tethers are associated with the positively and negatively curved membrane at the tether circumference and base, respectively. Thus a tempting hypothesis emerges that hypoxia-induced lipidome rewiring might affect lipid sorting leading to selective enrichment of inverted cone-shape lipids, that may foster the formation of highly curved regions at the site of cell membrane insertion, eventually leading to more frequent membrane nanotube generation.

### Hypoxia aggravates the migratory and invasive potential in pancreatic cancer

The aggressiveness of cancer cells is determined by their capacity to invade through the epithelial basement membrane. Hypoxia has been shown to promote cell migration and invasion in many cancers [46]. From a membrane biophysical perspective, the understanding of key determinants underlying tumor dissemination is limited, but changes in cell mechanics associated with lipidome and membrane remodelling are integral to metastatic dissemination. This is because cancer cells undergo extensive deformation to migrate through tissues and enter blood vessels [47, 48]. In accordance, recent studies have shown a strong correlation between decreasing cell stiffness and increasing invasive and metastatic efficiency [49-51]; the former seen in the hypoxic perturbation of cortical stiffness in a cell-line dependent fashion.

Using transwell migration and invasion assay, we observed that hypoxic pancreatic cancer cells exhibit enhanced migration and invasion compared to when in normoxic conditions (Fig. 5A-B). No change was seen in healthy pancreatic cells (Fig. S4A). Moreover, the migratory and invasive potential induced by hypoxia was more pronounced in PANC-1 cells compared to CAPAN-2. Accumulation of unsaturated fatty acids including PUFAs has been positively correlated with higher migration and invasion [52]; a higher increase of these species was seen in PANC-1 (Fig. 2A, C). Similarly, an increased abundance of cholesterol and cholesterol esters (CE) aggravates the metastatic potential in many cancers such as renal cell carcinoma, prostate, breast, and pancreatic cancer [53-56]. The accumulation of these in a cell-line-dependent fashion further corroborates a higher migratory and invasion potential in PANC-1 and CAPAN-2 cells induced under hypoxia. Finally, decreased cell stiffness in PANC-1 also correlates well with these observations. Surprisingly, enhanced metastasis index in endometrial cancer progresses through the coordinated action of the EMT pathway and higher cell membrane fluidity [52]. Our data suggest the interplay of additional factors in hypoxic pancreatic cancer cells without the involvement of cell membrane fluidity. Furthermore, the PM tension in PANC-1 was attenuated under hypoxia and supports a higher migratory potential, as increased PM tension is associated with inhibited cancer cell migration and invasion by modulating membrane curvature formation mediated by both lipids and membrane proteins [43]. Downregulated alpha-Linolenic acid and upregulated endocytosis pathway also might have contributed to such high migration and invasion potential [57, 58].

**Figure 5.**
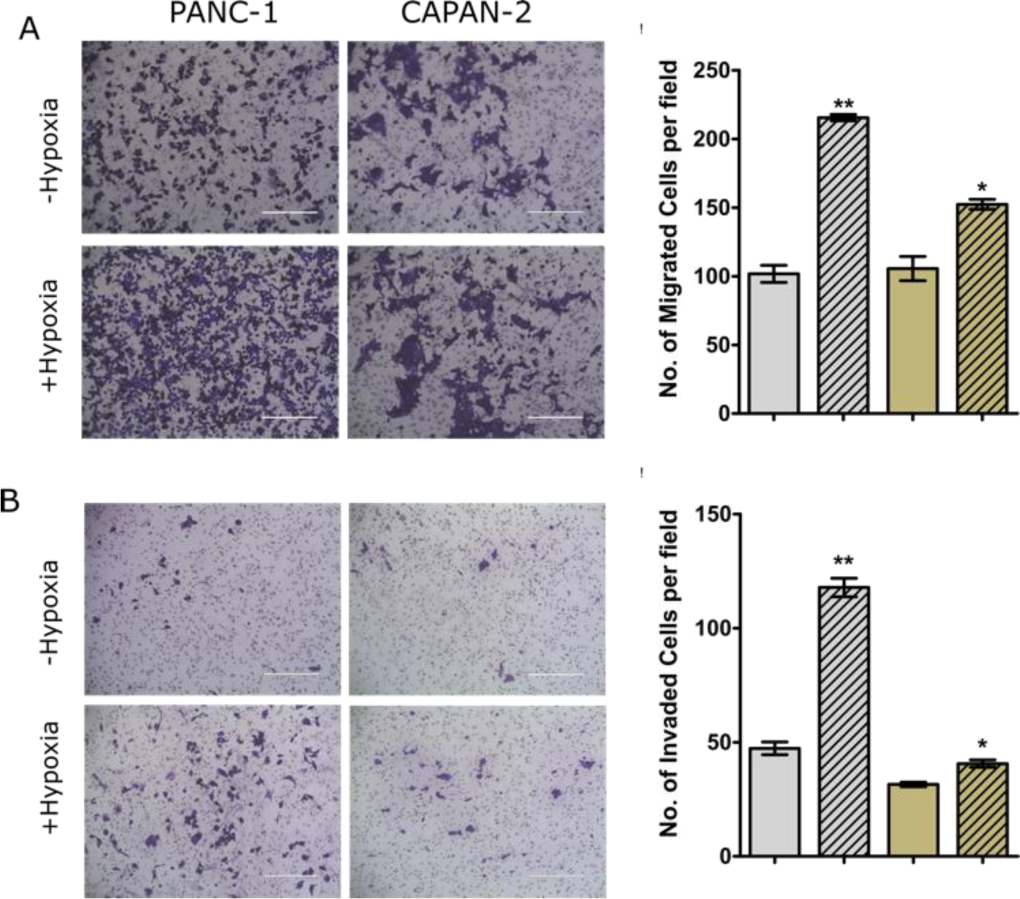
Hypoxia promotes migratory and invasive potential.

A. Hypoxia shows a higher cellular migration in transwell migration assay based on the visualization and quantification of the number of migratory cells compare to control cells was more.

B. Transwell matrigel invasion assay showed a significant increase in the invasive potential of hypoxia-treated PANC-1 and CAPAN-2 cells. The significance test was performed using a student’s *t*-test (\**P* < 0.01, two-tailed Student’s t-test).

In all the cases, bar plots with patterns represents hypoxic conditions. PANC-1, and CAPAN-2 represented by grey and brown colour codes, respectively.

## Discussion

Hypoxia plays an active role in promoting tumor survival, progression, and invasion and is a common phenomenon in most malignant tumors, especially in pancreatic cancer. Thus, studying the effect and response of hypoxic microenvironment in pancreatic cancer is gaining attention as an important therapeutic target. To adapt to the severe hypoxia environment, in general, cancer cells change their metabolic phenotypes to maintain their survival and proliferation. Among them, aberrant metabolism of fatty acids, and lipids are some of the well decorated hallmarks. The foremost impact of altered lipid metabolism is on the biophysical properties of cellular membranes, including plasma membrane. Cell membranes orchestrate various cellular functions by compartmentalizing proteins and lipids in distinct membrane domains, modulating protein structure and function, lipid and protein. In here, membrane biophysical properties such as membrane order, hydration, fluidity, curvature, tension etc all have specific roles in regulating biomolecule function by regulating their sorting, structure, diffusion and interactions with other partners. Collectively, an robust homeostatic response of mammalian membranes is crucial in both normal and disease states and led us to explore if and how the same is achieved and implemented under hypoxic environment in pancreatic cancer.

Lipidomic analysis of hypoxic pancreatic cancer cells revealed enhanced accumulation of TG and CE, while downregulation of GPLs and SPs Increased abundance of CE and TG underlined a significant induction of lipid droplets. Lipid species with longer acyl chains lengths and increased desaturation levels showed a significant increase under hypoxia, and was concomitant with an increase of lipids with saturated lipid chains. Furthermore, typical acyl-containing species were upregulated and alkyl- and alkenyl-containing “ether” PC, LPE and TG lipid species were downregulated and correlates well with the cellular abundance of specific enzymes such as AGPS also modulated under hypoxic stress. Biophysical membrane assays reported a tight maintenance of membrane properties such as fluidity and order, while the cell cortical stiffness decreased along with reduction of actin volume. In addition, facile tether dynamics was observed and could be critical for the observed enhanced migration and invasion under hypoxic states. In summary, we show a correlation or dependence of distinctive lipid remodelling upon hypoxia induction in pancreatic cancer and how the same is buffered enabling a homeostatic membrane fluidity response. The latter accounts for a robust migration and invasion phenotype in hypoxic pancreatic cancers. Deeper understanding of the molecular mechanisms behind this phenomena could furnish tangible avenues for therapeutic targeting of pancreatic cancer using membrane-centric interventions.

## Data availability

The data described in this article will be shared upon request. Inquiries should be directed to and will be fulfilled by the lead contact Shobhna Kapoor, Department of Chemistry, Indian Institute of Technology Bombay, India.

## Acknowledgments and Funding

This work was supported by the DBT/Welcome Trust India Alliance Fellowship (IA/I/21/1/505624) awarded to SK. This work is also supported by grants from DST-SERB (EMR/2016/005414, and WEA/2020/000032). Central Confocal and AFM Facilities at IIT Bombay are gratefully acknowledged. We are thankful to Prof. Ritu Aneja for the PANC-1 and CAPAN-2 cell lines.

