## Supplementary file 1 for "Lipid remodeling by hypoxia aggravates migratory potential in pancreatic cancer while maintaining membrane homeostasis"

This supplementary file contains SI methods, and 4 SI Figures.

### Materials and Methods:

#### Materials:

Foetal bovine serum (FBS, RM1112), Dulbecco's Modified Eagle Medium (DMEM-AT007), 1X Dulbecco's phosphate buffer saline (D-PBS), 20X Antibiotic-antimycotic solution, 0.25% trypsin-EDTA, MTT reagent and RIPA buffer were procured from HiMedia, India. Laurdan dye, D-5030 media, puromycin, hEGF, Cobalt (II) chloride hexahydrate (C8661) and Bradford reagent were purchased from Sigma Aldrich, St. Louis, Missouri, USA. The probes DPH (diphenylhexatriene) and TMA-DPH (trimethylamino-diphenylhexatriene) were purchased from Cayman Chemicals, USA. Di-4-ANEPPDHQ RDHPE TRITC-Phalloidin and Bodipy dyes were purchased from Thermo Fischer Scientific. Western blotting substrate and protease inhibitor tablets were also from Thermo Fischer Scientific, USA. PhosphoSTOP tablets were purchased from Roche, Switzerland. Mouse monoclonal anti-HIF1α antibody was purchased from BD Biosciences, San Jose, CA, Mouse monoclonal anti-β-actin antibody from LI-COR Biosciences and HRP-conjugated goat monoclonal anti-mouse secondary antibody was from Thermo Fischer Scientific. M3 base medium without antibiotic was purchased from INCELL Corporation, LLC.

**Cell culture:**

PANC-1 and CAPAN-2 pancreatic ductal adenocarcinoma (PDAC) cells were provided as a generous gift by Prof. Ritu Aneja from the Georgia State University, Atlanta. The normal HPNE cell line was purchased from ATCC. PANC-1 & CAPAN-2 cell lines were cultured in DMEM medium and HPNE cells were maintained in D5030 medium supplemented with M3 base medium.

**Cobalt Chloride ($Cocl_2$) Treatment to mimic hypoxia:**

$Cocl_2$ was dissolved in autoclaved MilliQ water and the stock was diluted in serum free medium (0.4%FBS) to achieve the required concentration.

**Cell viability assay:**

Cytotoxic effect of $Cocl_2$ was evaluated using MTT assay. Briefly, all the pancreatic cells were seeded in a flat bottomed 96-well polystyrene coated plate at a seeding density of $1*10^4$ cells in duplicates. After 36 hours cells were treated with different concentrations of $Cocl_2$ from 800 µM to 50 µM (2 fold dilutions) for 24 hours in high and low serum media. After treatment with each diluted compound and incubation for 24 hour, 100µL of MTT (0.5 mg/ ml) was added to each well and incubated for 3 hours at $37^oC$. Formazan crystals formed after 3 hours in each well were dissolved in 150 µL of DMSO and the plates were read immediately in a microplate reader (BIO-RAD microplate reader-550) at 570 nm. Wells without cells were used as blanks. This method was used to determine the cell viability.

LC-MS/MS data for lipidomic study was searched through MS Dial 4.90. The mass accuracy settings are 0.005 Da and 0.025 Da for MS1 and MS2. The minimum peak height is 5000 and mass slice width is 0.05 Da. The identification score cut off is 80%Post identification was done with a text file containing name and m/z of each standard (SPLASH and Deuterated Ceramide lipidomix Mass Spec Standard). In positive mode, [M+H]+, [M+NH4]+ and [M+H-H2O]+ were selected as ion forms. In negative mode, [M-H]- and [M+CH3COO]- were selected as ion forms. All lipid classes available were selected for the search. The retention time tolerance for alignment is 0.1 min. Lipids with maximum intensity less than 5-fold of average intensity in blank was removed. All other settings were default. All lipid LC-MS features were manually inspected and re-integrated when needed. These four types of lipids, 1) lipids with only sum composition except SM, 2) lipid identification due to peak tailing, 3) retention time outliner within each lipid class, 4) LPA and PA artifacts generated by in-source fragmentation of LPS and PS were also removed. The shorthand notation used for lipid classification and structural representation follows the nomenclature proposed previously (Liebisch G, Fahy E, Aoki J, Dennis EA, Durand T, Ejsing CS, et al. Update on LIPID MAPS classification, nomenclature, and shorthand notation for MS-derived lipid structures [2].

Label free proteomics data analysis by proteome discoverer 2.4:

The raw data sets were processed using Proteome Discoverer (PD; version 2.4. Using the parameters discussed in a previous study [5]. In brief, the variable modification of Carbamidomethyl and oxidation was set as static modification. The search engines used were MASCOT and SEQUEST with the Homo sapiens database. False-Discovery Rates (FDRs) for peptide and protein identifications were set to 1%. For data normalization the ‘Total peptide intensity’ parameter was used. Volcano plot was generated using the metaboanalyst 5.0 software and, significant down regulated and upregulated protein lists were collected for the further data annotation and interpretation.

**Steady-State Fluorescence Anisotropy:**

For the study of membrane fluidity by steady state fluorescence anisotropy, $3*10^5$ of each cells were seeded per well in 6 well plate and allowed to attach for 36 h. Cells were treated for 24 h with 200 µM $Cocl_2$ in low serum condition. Post treatment, cells were washed with 1X PBS, trypsinized, centrifuged for 5 min at 2500 rpm, and suspended in 1 mL of PBS. Laurdan, TMADPH and DPH probes were dissolved in DMSO and added to achieve a final probe concentration of 5 µM and 4 µM respectively. Cells were incubated at room temperature for 20 min, centrifuged at 2500 rpm for 10 min and re-suspended in PBS. Fluorescence anisotropy was measured using a Varian Cary fluorescence spectrophotometer with a temperature controller equipped with Varian manual polarizer accessory (Australia). The cell suspension after transferring to a cuvette kept sometime to attain temperature of 37°C using a circulating water bath. The samples were excited with vertically and horizontally polarized lights and respective polarized emission intensities were recorded. The degree of fluorescence steady-state anisotropy (r) was calculated from the following equation:

$$r = \frac{I_{VV} - G * I_{VH}}{I_{VV} + 2G * I_{VH}}$$

where IVV and IVH are the vertically polarized emission intensities measured in directions parallel (both at 0°, $I_0$) and perpendicular (excitation: 0°, emission: 90°, $I_{90}$) to the excitation beam. The correction factor (G= IHV/IHH) is the ratio of the detection system sensitivity for the polarized light, where IHV and IHH corresponds to the emission intensities perpendicular and parallel to the horizontally excitation light. The correction due to cell scattering was applied for obtaining the final anisotropy values [6].

**FliptR FLIM imaging of pancreatic cancer cells:**

The FliptR probe includes two fluorophores which can twist between a planar (red) state under high tension and a twisted (green) state under lower tension. The carbon bond around which the fluorescent groups (green) can twist is shown in red. In more ordered phases, lipid packing is higher, as acyl chains apply a higher pressure that planarizes the probe and this leads to changes in excitation maxima with a red shift of 50–90 nm and thus an increase in fluorescence lifetime. This dye reports membrane tension. FLIM imaging was performed using a Nikon Eclipse Ti A1R microscope equipped with a time-correlated single-photon counting module from PicoQuant58. Excitation was performed using a pulsed 485 nm laser operating at 20 MHz, and the emission signal was collected through a 600/50 nm bandpass filter using a gated PMA hybrid 40 detector and a TimeHarp 260 PICO board (PicoQuant). SymPhoTime 64 software (PicoQuant) was then used to fit fluorescence decay data (from full images or regions of interest) to a dual exponential model [9].

Half life ($t_{1/2}$)= $\frac{ln2}{k}$

Diffusion coefficient (*D*) = $\frac{\varpi^2}{4*t_{\frac{1}{2}}}$

% mobile fraction: $\frac{I_{\infty}-\ I_a}{1-I_a}$

Where, I∞ is the full-scale normalized fluorescence intensity after full recovery and $I$α the normalized intensity of the first post-bleach.

**Lipid Droplet staining with Bodipy:**

After the hypoxia induction for 24h pancreatic cells were fixed with 4% paraformaldehyde for 20 min at RT and rinsed with 1X PBS for three times. Cells were then incubated with 1mM of BODIPY for 15 min in the dark, washed 2x with 1 PBS, incubated with 1 mg/mL DAPI for 15 min in the dark and washed 2x again with PBS. Coverslips were then mounted on glass slides and Z stack images were captured using confocal laser scanning microscope and processed with ImageJ software.

**Transwell migration and invasion assays:**

Transwell migration and invasion assays were performed to test migratory and invasive potential of pancreatic cancer cells upon hypoxia induction. Briefly, 4×104 cells were seeded in Boyden chambers (PET membrane, 8 µM pore size) with or without matrigel coating (Corning), in 0.4% serum media, in presence or absence of $CoCl_2$. The Boyden chambers were then inserted into 24-well plates containing DMEM supplemented with 10% fetal bovine serum as chemoattractant. Cells that migrated across the membrane were imaged 24 h post treatment. Before imaging, chambers were washed in PBS, fixed in methanol, followed by staining with a 0.5% crystal violet solution. The membranes were then imaged at 4 randomly selected fields with an inverted light microscope and cells no was calculated per field of the image.

**Statistical analysis:**

The results of independent experiments are presented as mean or median values as indicated. Error bars represents standard error of the mean (SEM). Statistical significance was tested using unpaired two-tailed t-test or two way ANOVA followed by Bonferroni's multiple comparison test as indicated. For AFM results non-parametric unpaired Mann-Whitney test was used. *P*-values < 0.05 were considered statistically significant.

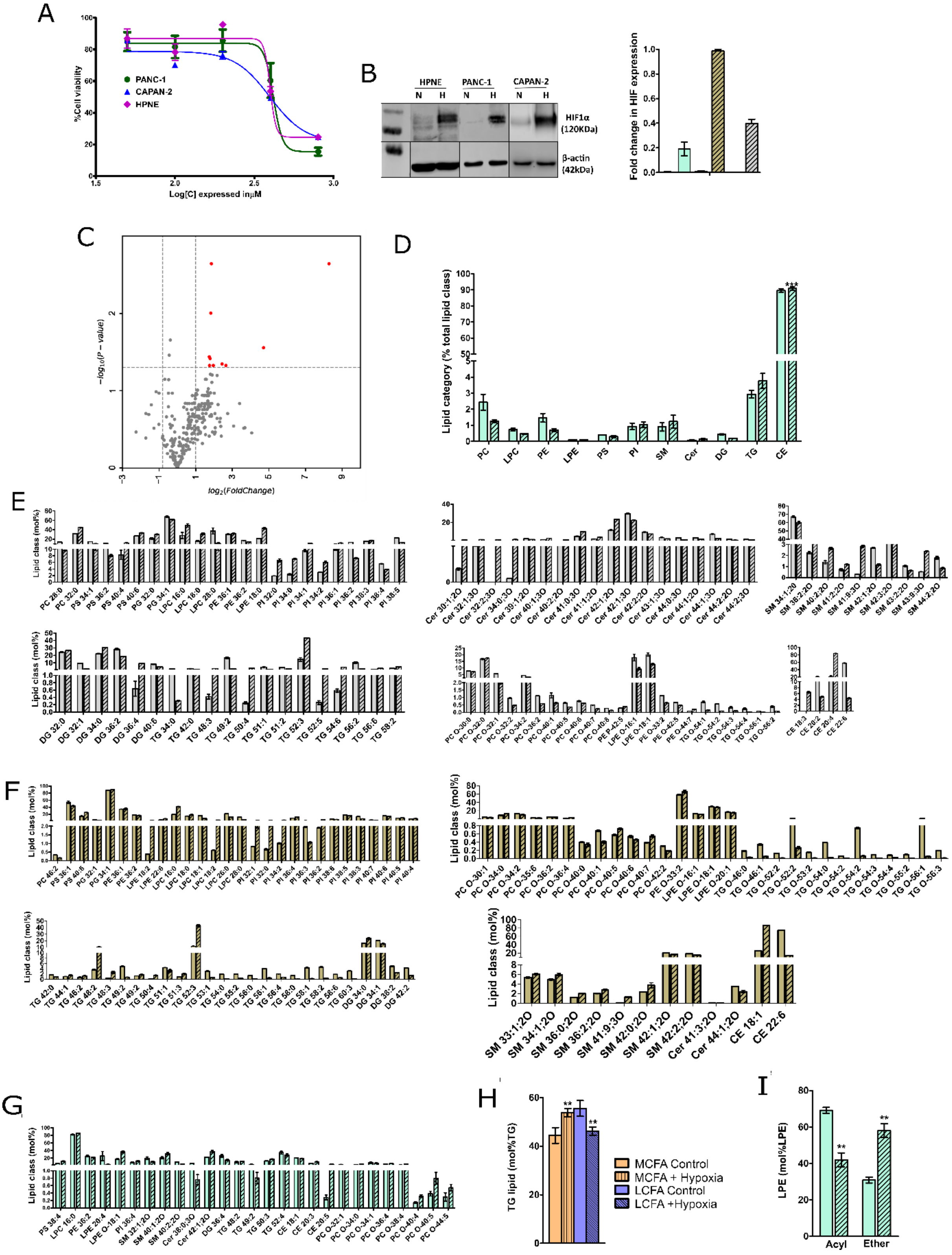

Figure S1. Lipidome remodeling induced by hypoxia.

A. Cell viability was determined using MTT assay. Cells were treated with different concentrations of $CoCl_2$ (50/100/200/400/800 µM) for 24h in a total of 3 replicates to select a concentration (200 µM) below $IC_{50}$ to be used for further experiments.

B. Cells treated with 200 µM of $CoCl_2$ showed a higher induction of HIF1-α. Quantification of the Western blot result was performed by calculating the ratio of the HIF-1α and β-actin and then the fold change in treatment over the control was plotted.

C. Volcano plot showing change in lipidome modulation of HPNE cells cultured in absence and presence of hypoxia.

D. Mol% total lipid abundance at the lipid category level of HPNE treated cells compare to control cells shows no significant change.

E, F & G. Mol% total significant lipid class abundance distribution of PANC-1, CAPAN-2 & HPNE respectively.

H & I. Mol% total lipid class abundance distribution at the lipid class level of HPNE as per chain length and acyl and ether lipid.

In all the cases, bar plots with patterns represents hypoxic conditions. PANC-1, CAPAN-2 and HPNE are represented by grey, brown and light green colour codes, respectively.

.

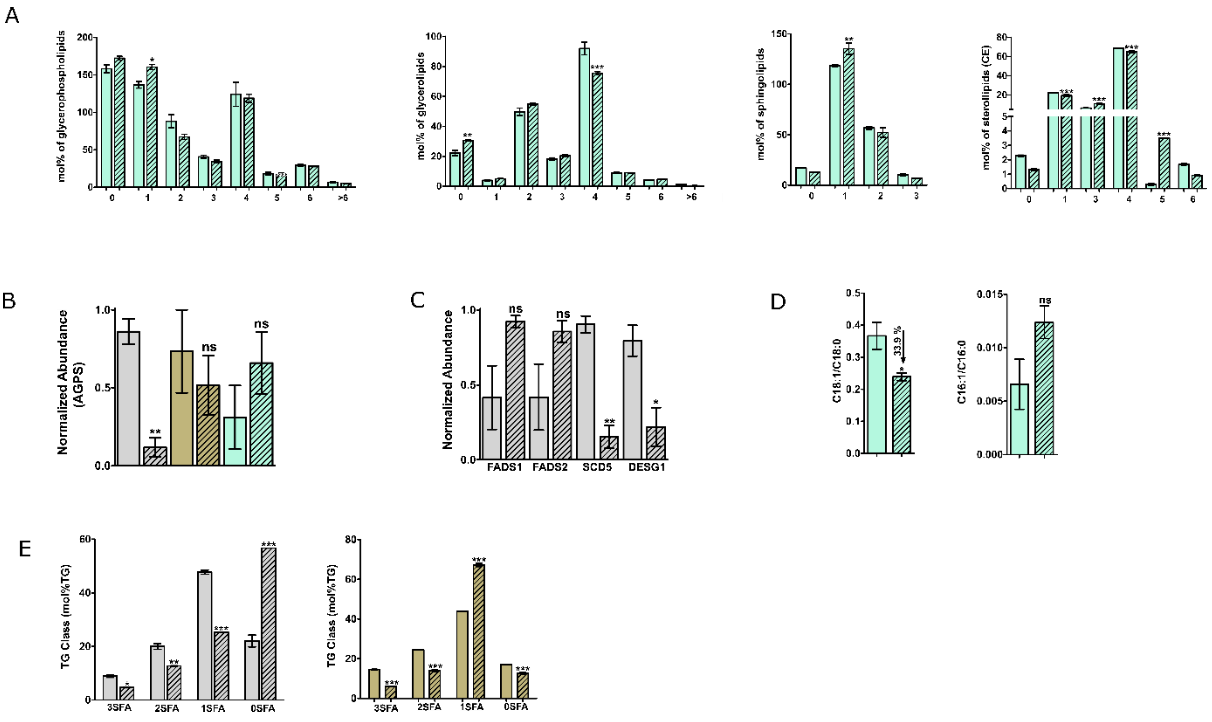


Figure S2. A. Mol% total lipid abundance at the lipid category and class level of HPNE showing different level of distributions of four lipid categories, including GPLs, SPs, GLs and sterol lipids ST.

B & C. Normalized abundance of AGPS, FADS1, FADS2, SCD5 and DESG protein in presence and absence of hypoxia.

D. Desaturation index (C18:1/C18:0) & (C16:1/C18:0) of LPC in HPNE.

E. Distribution of saturated and unsaturated TG species of PANC-1 and CAPAN-2 cells.

In all the cases, bar plots with patterns represents hypoxic conditions. PANC-1, CAPAN-2 and HPNE are represented by grey, brown and light green colour codes, respectively..

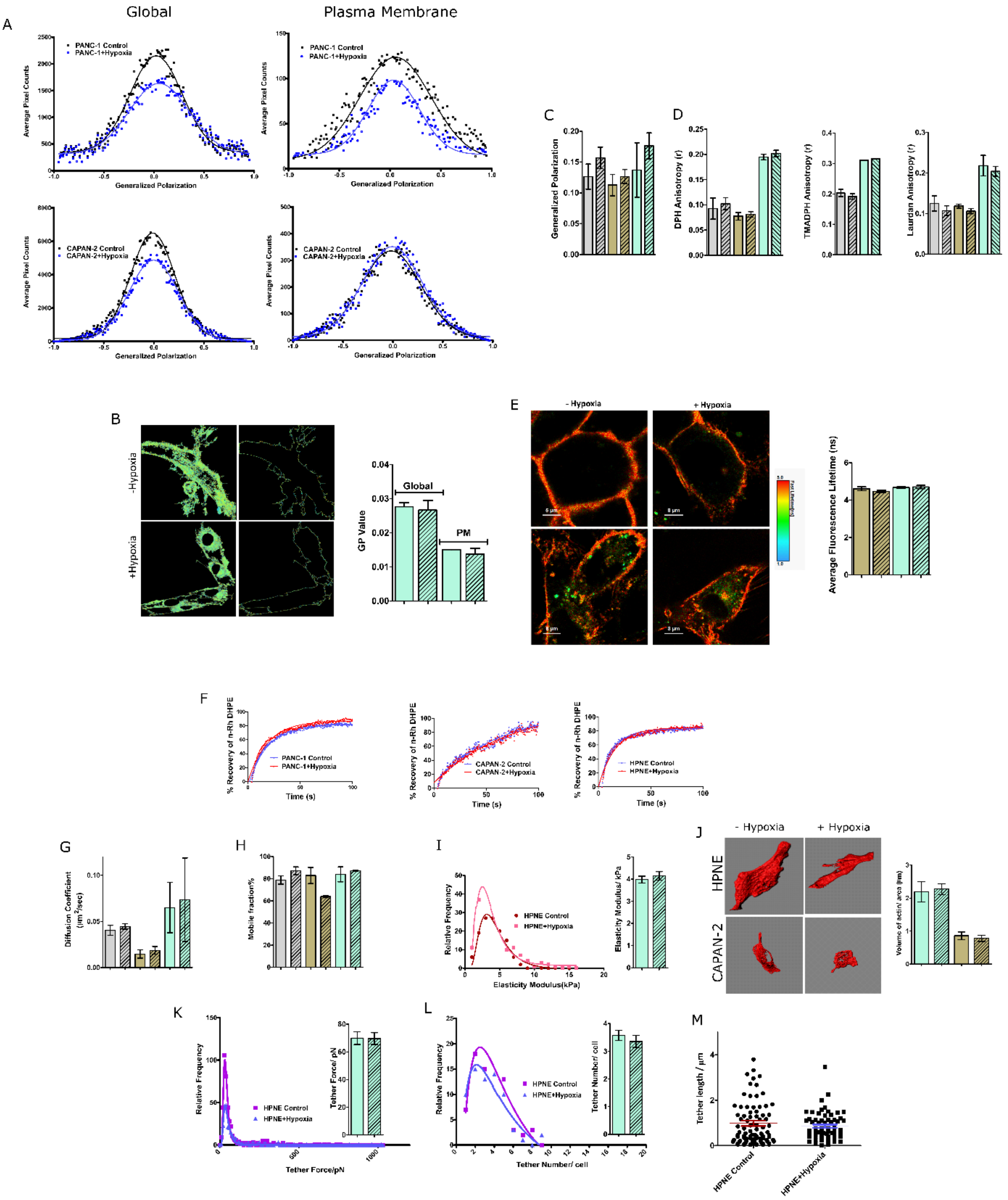


Figure S3. Membrane homeostasis remain intact in presence of hypoxia.

A. Global and segmented plasma membrane GP distribution from the stack of GP images (n = 85, N = 3) deconvoluted by fitting Gaussian distributions.

B. GP imaging of di-4-ANEPPDHQ-labeled cell membranes of HPNE cell lines shows no change in membrane packing after 24 h of hypoxia induction.

C. Laurdan spectroscopic data collected from control and hypoxia induced cells of three replicates were plotted to get generalized polarization value.

D. Pancreatic cells in the presence and absence of hypoxia were used to measure DPH, TMADPH and Laurdan fluorescence anisotropy of three independent experiments. Both the spectroscopy and anisotropy data showed non-significant change after statistical analysis was determined using student's t-test (*P < 0.01, two-tailed Student's t-test).

E. FLIM imaging of CAPAN-2 and HPNE cells with FliptR dye showed no significant change in average fluorescence lifetime after the 24h of hypoxia induction.

F. Normalized fluorescence recovery curves in presence and absence of hypoxia.

G & H. Bar graph showing the diffusion coefficients and the % mobile fraction of nRh-DHPE calculated as per the formula mentioned from the individual recovery curves (n>30 cells per group). Data were represented as the means ± SEM of three independent experiments and the statistical analysis was determined using student's t-test.

I. Elastic moduli distribution of HPNE cells in presence and absence of hypoxia. No significant change in elastic moduli is observed.

J. Representative isosurface images for HPNE & CAPAN-2 cells were used to quantify the actin abundance. Bar graph represents the volume of actin per unit area in control and hypoxia induced cells. Scale bar, 10 µm; 63X oil objective.

K. Relative frequency distributions of membrane tether forces, L. tether number and M. tether length in presence and absence of hypoxia. The mean values are provided in the bar graph. Significance test was performed using student's t-test (*P < 0.01, two-tailed Student's t-test). All the three relative frequency distributions were found to be non-significant after the student's t-test statistical analysis.

In all the cases, bar plots with patterns represents hypoxic conditions. PANC-1, CAPAN-2 and HPNE are represented by grey, brown and light green colour codes, respectively.

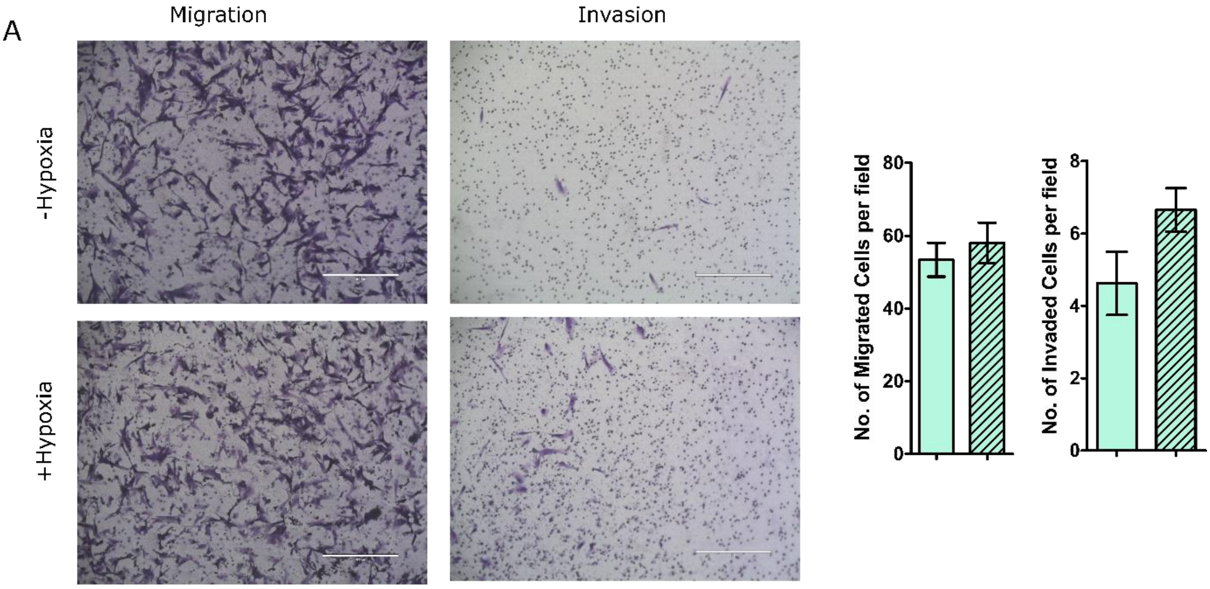


Figure S4. A. Hypoxia shows no change in cellular migration and B. invasion in HPNE cells. Significance test was performed using student's t-test (*P < 0.01, two-tailed Student's t-test). In all the cases, bar plots with patterns represents hypoxic conditions. PANC-1, CAPAN-2 and HPNE are represented by grey, brown and light green colour codes, respectively.
